# Local regulation of lipid synthesis controls ER sheet insertion into nuclear envelope holes to complete nuclear closure

**DOI:** 10.1101/757013

**Authors:** Lauren Penfield, Raakhee Shankar, Erik Szentgyörgyi, Alyssa Laffitte, Michael Mauro, Anjon Audhya, Thomas Müller-Reichert, Shirin Bahmanyar

## Abstract

The nuclear permeability barrier depends on closure of holes in the nuclear envelope (NE). Here, we use meiotic *C. elegans* oocytes to demonstrate that local control of glycerophospholipid synthesis by CNEP-1/CTDNEP1 regulates the insertion of ER sheets into NE holes and functions independently of ESCRT-III to ensure NE closure. Deletion of CNEP-1 causes excess incorporation of ER membranes into NE holes and a defective NE permeability barrier. ESCRT-III components accumulate at the NE opening surrounding the meiotic spindle, and loss of NE adaptors for ESCRT-III exacerbates NE sealing defects in *cnep-1* mutants. Limiting ER sheet production by restoring glycerophospholipid synthesis in *cnep-1* mutants rescued NE permeability defects. 3D analysis showed that membrane sheets feed into and narrow NE holes occluded by meiotic spindle microtubules supporting a role for ER sheet insertion in NE closure. Thus, feeding of ER sheets into NE holes must be coordinated with production of ER sheets near the NE to promote NE closure.

## Introduction

The nuclear envelope (NE) is a sealed spherical sheet that surrounds and protects the genome. The NE is composed of two continuous lipid bilayers (the inner and outer nuclear membranes) separated by a single lumen. The membranes and lumen of the NE are contiguous with the endoplasmic reticulum (ER), a large interconnected membrane network that extends throughout the cytoplasm. Lipid metabolizing enzymes associated with ER membranes produce the bilayer glycerophospholipids of the NE (Hetzer, 2010; Baumann and Walz, 2001; Fagone and Jackowski, 2009). A subset of integral membrane proteins synthesized by ER-associated ribosomes are enriched at the inner nuclear membrane (INM), giving the NE its unique identity. In open mitosis, ER membranes and associated INM proteins wrap the surface of segregated chromosome to initiate the formation of the NE. Completion of the NE requires the closure of holes that coincide with spindle microtubules, as well as insertion of nuclear pore complexes (NPCs) for transport between the nucleus and cytoplasm (Hetzer, 2010; Ungricht and Kutay, 2017).

The NE must be sealed at the end of mitosis to serve its selective permeability barrier function. Assembly of the cytoplasmic protein complexes of ESCRT-III on the negatively curved surface of NE holes constricts membranes to execute fission and seal the NE at the end of mitosis (Olmos et al., 2015; Vietri et al., 2015). The ESCRT-II/III hybrid protein CHMP7 and the INM protein LEM2/LEM-2 form a complex at NE holes to recruit ESCRT-III components (Olmos et al., 2016; Gu et al., 2017; Webster et al., 2016). These components in turn recruit the microtubule-severing AAA-ATPase, spastin, to coordinate mitotic spindle disassembly with membrane fission (Vietri et al., 2015). This sealing process restricts traffic to the NPCs through which molecules greater than ∼40 kDa cannot diffuse passively (Ungricht and Kutay, 2017; Wente and Rout, 2010). Without ESCRT-mediated NE sealing, ∼40-60 nm holes occupied by spindle microtubules persist and cause slow nucleocytoplasmic mixing, abnormal nuclear morphologies and DNA damage (Olmos et al., 2015; Vietri et al., 2015; Olmos et al., 2016; Ventimiglia et al., 2018).

In interphase, NE ruptures that cause rapid mixing of nuclear and cytoplasmic components, and at times nuclear entry of whole organelles, depend on ESCRT-III for resealing (De vos et al., 2011; Vargas et al., 2012; Denais et al., 2016; Raab et al., 2016). Even micron-scale punctures caused by lasers undergo slow repair (Denais et al., 2016; Penfield et al., 2018; Halfmann et al., 2019). Given the evidence that the ESCRT-III spiral filament (VPS-32/Snf7/CHMP4B) resolves holes of ∼30-50 nm in diameter (Wollert and Hurley, 2010; McCullough et al., 2018; Olmos et al., 2015; Olmos and Carlton, 2016), mechanisms must exist to narrow large membrane holes in interphase NE ruptures and during NE reformation before ESCRT-III acts. ESCRT-machinery may have a direct role in the constriction of larger holes. In fact, ESCRT-III components line the >1 μm wide cytokinetic bridge (Elia et al., 2011; Guizetti et al., 2011), although exactly how constriction occurs at the site of abscission remains unclear (McCullough et al., 2018; Henne et al., 2013). The NE-specific adaptors for ESCRTs, CHMP7 and LEM2/LEM-2, may contribute to constricting large NE openings, although currently evidence for this does not exist.

Contiguous ER membranes are a potential source of membranes that narrow large NE openings prior to closure by ESCRT-III. In the rapid cell cycles of *Drosophila* embryos, specialized ER membrane sheets containing pre-assembled NPCs insert into NE openings to facilitate nuclear expansion during interphase (Hampoelz et al., 2016). However, the source of the membrane to close NE openings at the end of mitosis or after rupture is not known.

From yeast to metazoans, local regulation of glycerophospholipid synthesis limits incorporation of membranes into the NE, and lipin is the key enzyme controlling the production of glycerophospholipids in the ER/NE network (Siniossoglou, 2013; Bahmanyar, 2015). Lowering of lipin activity causes NE expansion in budding yeast and abnormal nuclear morphologies after open mitosis in *C. elegans* (Santos-Rosa et al., 2005; Golden et al., 2009; Gorjánácz and Mattaj, 2009; Bahmanyar et al., 2014). Lipin dephosphorylates phosphatidic acid to produce diacylglycerol that is used for the synthesis of the major structural glycerophospholipids, phosphatidylcholine (PC) and phosphatidylethanolamine (PE), at the expense of the synthesis of phosphatidylinositol (PI) (Han et al., 2006; Siniossoglou, 2013; Bahmanyar, 2015). Lipin is inhibited by multi-site phosphorylation and activated by dephosphorylation by the integral membrane phosphatase Nem1/CNEP-1/CTDNEP1 of the NE (Santos-Rosa et al., 2005; O’Hara et al., 2006; Han et al., 2012; Kim et al., 2007; Peterson et al., 2011; Grimsey et al., 2008). In *C. elegans* embryos, CNEP-1 locally activates lipin to bias the flux of glycerophospholipids away from PI synthesis and restrict ER sheet formation proximal to the NE (Bahmanyar et al., 2014). Deletion of CNEP-1 from embryos increased PI levels and caused formation of ectopic ER sheets close to the NE that wrap around the permeabilized NE upon entry into mitosis (Bahmanyar et al., 2014). Whether local activation of lipin by CTDNEP1/CNEP-1 is involved in closure of NE openings during NE formation had not been tested.

We report a role for CNEP-1 in biasing flux away from PI synthesis to promote closure of the NE in the *C. elegans* zygote. Loss of CNEP-1 disrupts NE closure around the acentriolar meiotic spindle by producing excess membranes that overflow into NE openings. We further show that loss of the NE-adaptors for ESCRT-III, LEM-2 or CHMP-7, exacerbates nuclear permeability barrier defects in *cnep-1* mutant embryos, suggesting CNEP-1 and the NE adaptors for ESCRT-III have partially overlapping roles in NE closure. Three dimensional reconstructions from electron tomograms of wild type meiotic oocytes revealed cytoplasmic membranes that feed into and narrow NE openings containing microtubules during NE formation, further suggesting that insertion of ER sheets into NE holes is a way to prevent macromolecules from freely traversing the NE. Our data define a previously uncharacterized role for ER membrane insertion in closure of NE openings and reveal that local glycerophospholipid flux is a mechanism that regulates the formation of the nuclear permeability barrier.

## Results and Discussion

We analyzed the role of NE adaptors for ESCRT-III in *C. elegans* during NE closure by monitoring the highly asymmetric closure of the NE around the acentriolar microtubule spindle during the final stages of oocyte meiosis II. Prior to fertilization oocytes are arrested in prophase of meiosis I (Oegema and Hyman, 2006). Fertilization by haploid sperm triggers oocytes to undergo two rounds of meiotic chromosome segregation. An acentriolar meiotic spindle surrounds sister chromatids in metaphase II (Albertson and Thomson, 1993; Oegema and Hyman, 2006; Fabritius et al., 2011) (Fig. 1 A) and transitions into a central spindle of microtubules between the segregated chromosomes in anaphase II (“maximal spindle shortening”, Fig. 1 A) (Yang et al., 2003; Fabritius et al., 2011; Redemann et al., 2018). As one set of segregated chromosomes extrudes to generate a second polar body, the haploid, oocyte-derived pronucleus forms. The spindle of microtubules elongate and dissipate as the oocyte-derived pronucleus forms (Yang et al., 2003; Fabritius et al., 2011; Redemann et al., 2018), but the mechanisms of remodeling and sealing the NE were not known.

**Figure 1.**
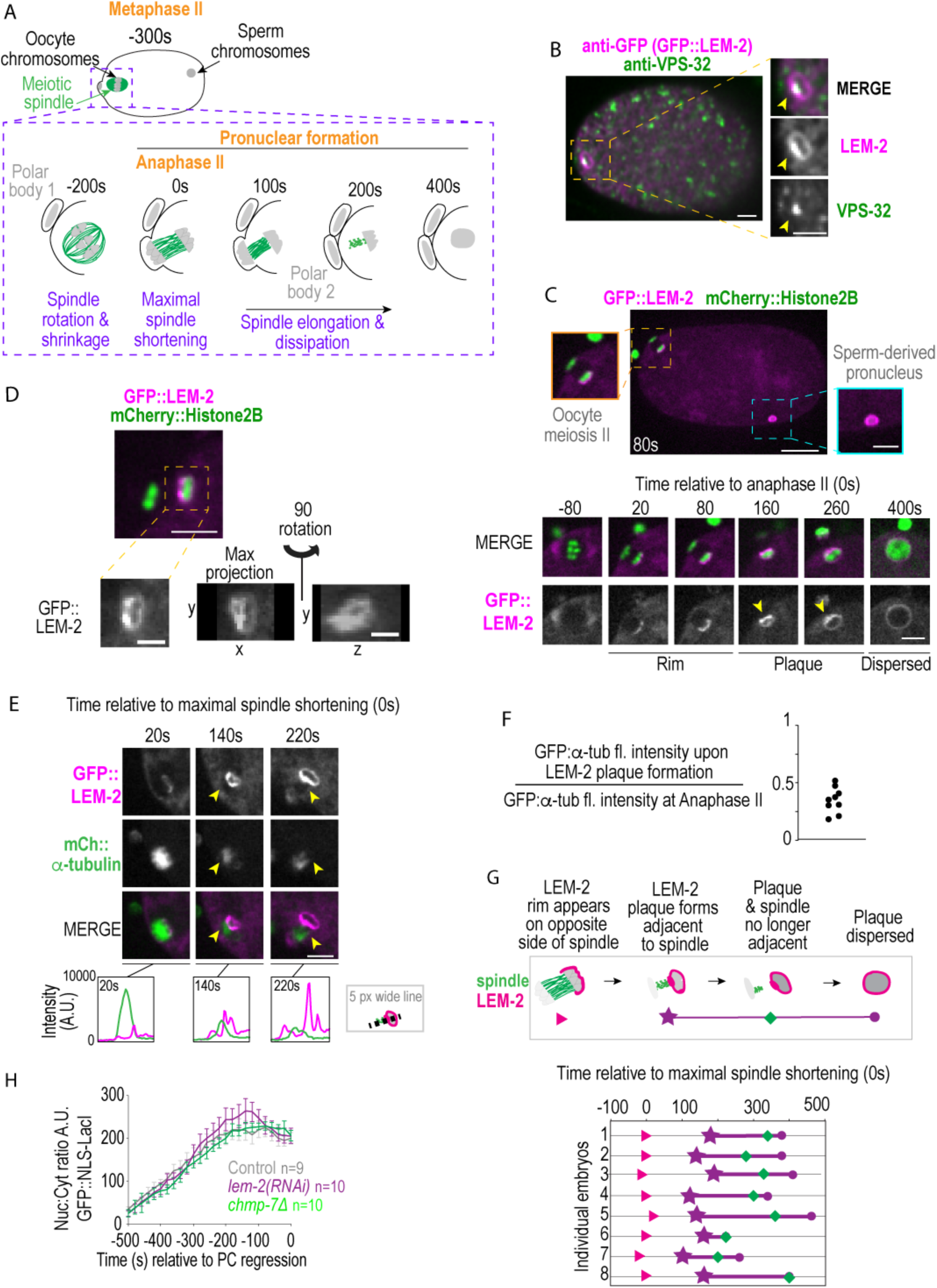
The nuclear envelope adaptor for ESCRT-III LEM-2 transiently accumulates to the site of nuclear closure that surrounds the disspitating meiotic spindle. **(A)** Schematic of events during and following *C. elegans* oocyte meiosis II. **(B)** Fixed overview and magnified images of *C. elegans* oocyte expressing GFP::LEM-2 immunostained for GFP and VPS-32. Arrowheads mark accumulation of LEM-2 and VPS-32 on the NE. Scale bars, 5 μm **(C)** Confocal image of embryo expressing GFP::LEM-2 and mCherry::Histone2B from a time-lapse series. Scale bar, 10 μm. Magnified images on left and right show sperm-derived-and oocyte-derived pronuclei. Below, confocal images of time lapse series of same oocyte-derived pronucleus. Scale bar, 5 μm. **(D)** Confocal image of *C. elegans* oocyte expressing mCherry::Histone2B and GFP::LEM-2. Scale bar, 10 μm. Below, magnified images of GFP::LEM-2 in a single z-slice, a max projection, and a 90° rotation. Scale bar, 2.5 μm. **(E)** Confocal images from time series of pronucleus of oocyte expressing GFP::LEM-2 and mCherry::α-tubulin. Below, line scans measuring the background-corrected fluorescent intensities. **(F)** Individual measurements of maximum fluorescence intensity of meiotic spindle at the time of appearance of a plaque of LEM-2. Time in seconds relative to the maximum fluorescence intensity. **(G)** Timeline of events scored from confocal time lapse series of individual oocytes expressing GFP::LEM-2 and mCherry::α-tubulin. Scale bars, 5 μm. **(H)** Plot of nuclear:cytoplasmic ratio of GFP::NLS-LacI for the indicated conditions. Average +/- SEM is shown. All measurements were corrected for background and normalized to nuclear area. n = number of embryos.

Fluorescence microscopy of fixed cells showed that the ESCRT-III adapter LEM-2 tagged with GFP marks the nuclear rim and forms a “plaque” containing ESCRT-III spiral component VPS-32 (CHMP4B) on the forming oocyte-derived pronucleus (Fig. 1 B). Live imaging revealed that GFP::LEM-2 initially appears at anaphase II on the chromatin farthest from the extruding polar body (Fig. 1 C and Fig. S1 A, 28 +/- 14 s, average +/- SD relative to anaphase II onset, n = 9). About 180 s after anaphase II onset, GFP::LEM-2 accumulates into a plaque opposite the initial rim (Fig. 1 C and Fig. S1 A, 182 +/- 32, n = 9) and then disperses ∼200 s later into a uniform rim around the expanding oocyte-derived pronucleus (Fig. 1 C and Fig. S1 A-C, 197 +/- 74 s, average duration +/- SD of plaque, n = 7). The general ER membrane marker (SP12::GFP) did not form a plaque on the oocyte pronucleus (Fig. S1 D), and LEM-2 did not form a plaque on the sperm-derived pronucleus (Fig. 1 C). Thus, ESCRTs specifically localize to a site that requires remodeling of the NE due to obstruction by meiotic spindle microtubules.

We hypothesized that the transient LEM-2 plaque accumulates at an opening of the NE surrounding the meiotic spindle and its dynamics may be correlated with spindle remodeling and disassembly. In 20% of the cases (n = 20 oocyte-derived pronuclei), GFP::LEM-2 puncta surrounded an ∼1 µm wide opening (Fig. 1 D, max projection), an observation that was also occasionally made for large openings caused by NE rupture (Fig. S1 E). In contrast to NE rupture sites, the large opening in meiosis is occupied by the meiotic spindle (Fig. S1 F). We reasoned that the LEM-2 plaque that we observed in other examples of meiotic NE formation corresponds to an opening in the NE. Live cell imaging showed that the fluorescence intensity of the GFP::LEM-2 plaque increases as the meiotic spindle microtubule density marked with mCherry::α-tubulin decreases (Fig. 1 E, compare 140 s to 220 s; Fig. 1 F, Video 1). This suggests that membranes accumulate LEM-2 as they coat the surface of chromatin within the region contacting the spindle, similar to human LEM2 in mitosis (Gu et al., 2017). The time course of LEM-2 plaque formation and dispersal correlates with spindle dynamics across many embryos (Fig. 1 G).

Although ESCRT-III machinery localizes to the ∼1 µm opening around the meiotic spindle, RNAi-depletion of LEM-2 or deletion of CHMP7 (*chmp-7*) (Fig. S1 G) did not impact the nuclear:cytoplasmic ratio of a GFP::NLS reporter (Fig. 1 H, GFP::NLS-LacI), so ESCRT-III is not required for narrowing of the large hole. In mammalian cells, loss of ESCRT-III machinery does not cause a substantial loss of a GFP::NLS reporter, however, the lack of sealing smaller holes around single microtubules causes slow mixing of nuclear and cytoplasmic contents and is detected in the next interphase (Olmos et al., 2015; Vietri et al., 2015; Olmos et al., 2016). Perhaps the short time (∼20 min) between the end of meiosis II and entry into mitosis in early *C. elegans* embryos may not be sufficient to attain measurable loss of imported proteins from small holes in the NE. Alternatively or in addition, there may be a redundant pathway to close NE openings and restrict nucleocytoplasmic exchange in *C. elegans* oocytes.

Three dimensional reconstructions from serial-section electron tomograms of early and late stage meiotic *C. elegans* oocytes showed that connected as well as discontinuous membranes fill in the large gap facing the meiotic spindle and transition into a continuous rim that contains small openings that intersect with a few remaining microtubules. In the early stage example, the large gap in the region of the NE facing the meiotic spindle (Fig. S1 H, 0.85 μm x 1.5 μm in this example) contains a segment with a few discontinuous membranes that coincide with microtubules (Fig. 2 A and Videos 2 and 3). Another segment of the gap contains a more continuous yet highly fenestrated nuclear rim (Fig. S1 H). Nuclear membranes in this region have clear connections with flat membrane sheets that are likely the ER (Fig. S1 H). The NE is mostly continuous on the side of chromatin opposite the spindle with the exception of some small holes had a diameter of 83 +/- 41 nm (average +/-SD, n = 33 holes; Fig. S1 H and Fig. 2 A, Videos 2 and 3), consistent with the nuclear membrane rim first observed 20 s after the onset of anaphase II (Fig. 1 C and Fig. S1 A). At a later time point, the nuclear membranes facing the meiotic spindle become more continuous (largest discontinuous region ∼400 nm in diameter) and contain a few remaining spindle microtubules that occupy holes with an average diameter of 133 nm (60 nm to 320 nm; n = 8 gaps, Fig. 2 B and C and Videos 4 and 5).

**Figure 2.**
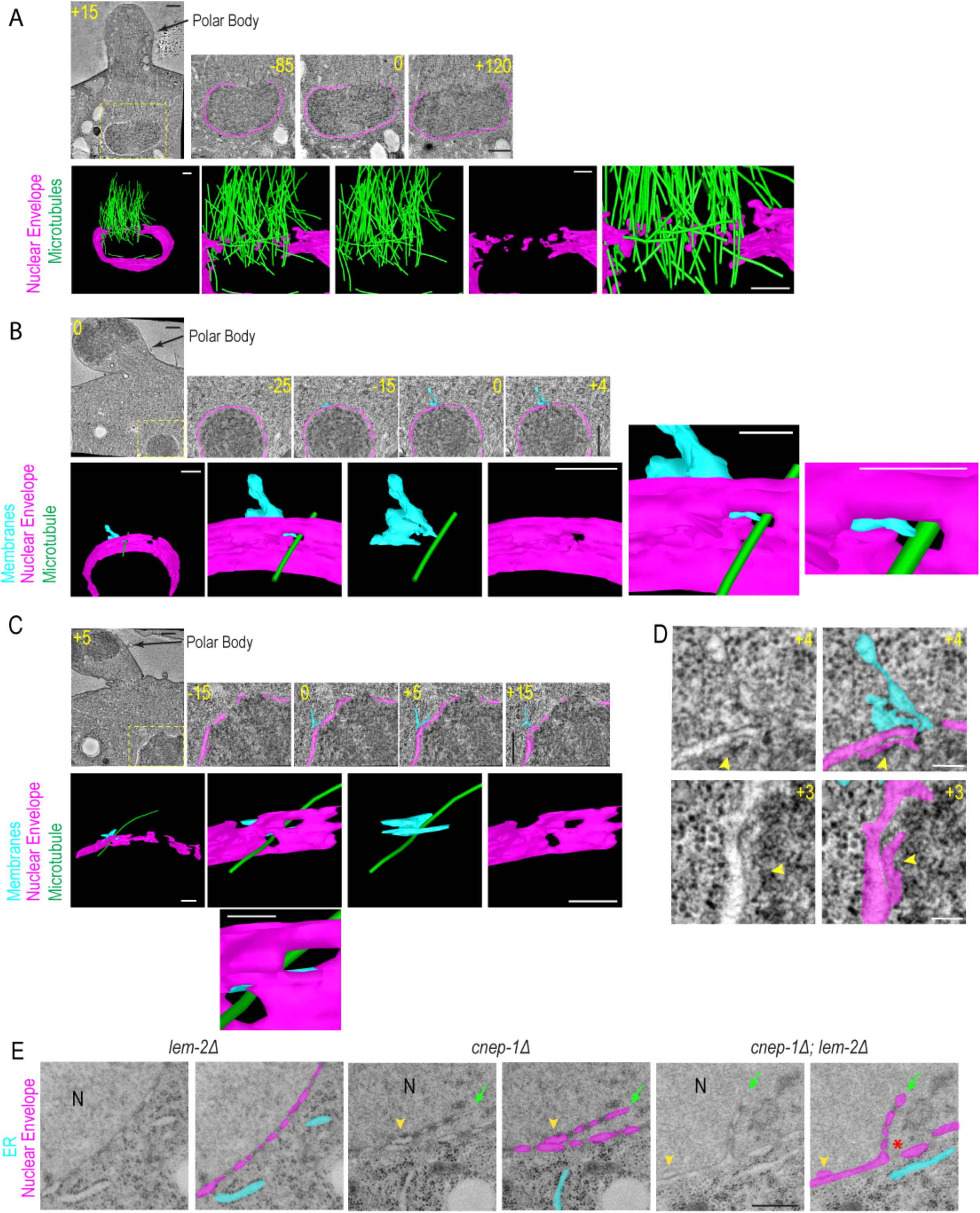
Membrane sheets feed into nuclear envelope holes that coincide with meiotic spindle microtubules. **(A-C)** Select images from electron tomograms of oocyte-derived pronucleus at mid-anaphase (A) and late-anaphase II (B-C). Left, overview image. Right, magnified z-slices of pronucleus (number indicates z-slice relative to indicated position 0) where the nuclear envelope (magenta) and contacting membranes (cyan) are traced. Scale bars, 500 nm. Below, 3D models generated from tracing the nuclear envelope (magenta), microtubules (green) and membrane sheets contacting the nuclear envelope (cyan). Scale bars, 250 nm. Scale bar for magnified image in (B), 100 nm. **(D)** Magnified images from traced and untraced electron tomograms in regions shown in B (left) and C (right). Arrowheads mark parallel membrane sheets in the nuclear interior. Scale bar, 100 nm. **(E)** Representative transmission electron micrographs for the indicated conditions. For each condition the untraced and traced image is shown. N notes the nuclear interior, arrowhead marks 2 parallel lumens, arrow marks NE extension. Scale bar, 500 nm.

At the later time points 17% of NE holes were directly adjacent to a membrane sheet that formed contacts with the outer nuclear membrane (n = 41 holes, Fig. 2 B and C). Of these holes, 71% (5/7) were occupied by microtubules. In one example, a flattened portion of a membrane sheet partially filled ∼50 nm of the ∼80 nm NE gap narrowing it to a size slightly larger than a single microtubule (Fig. 2 B and Videos 4 and 5). In a different example, a flattened region of a membrane sheet stretches and partially invades two large gaps (92 nm and 80 nm in the longest dimension), one that is occupied by a microtubule and one that is not (Fig. 2 C). Interestingly, this membrane sheet is also in contact with the cytoplasmic portion of the microtubule that occupies the NE opening (Fig. 2 C). Thus, membrane sheets, potentially directed by microtubules, insert into and narrow NE openings during the late stage of nuclear closure. Nearby 48% of the NE holes (n = 41), we also observed an additional extra parallel sheet that lay next to a region of the INM (Fig. 2 D). We predict that the redundant parallel membrane sheets (Fig. 2 D) are nuclear membranes displaced by incoming ER sheets that narrow NE openings.

The tomograms and our previous observation of parallel ER membrane sheets proximal to NE openings in CNEP-1 mutant embryos during mitosis (Bahmanyar et al. 2014) suggested that CNEP-1 may be involved in incorporation of ER membranes at NE holes. To test this idea, we analyzed TEM of thin sections of interphase nuclei of *C. elegans* embryos in *cnep-1Δ* and *lem-2Δ* mutants alone and *cnep-1Δ;lem-2Δ* double mutants (Fig. 2 E and Fig. S2 A-D). Interphase NEs in *cnep-1Δ* and *cnep-1Δ;lem-2Δ* double mutants contained NE regions in which two lumens could be traced (Fig. 2 E and Fig. S2 C-E; 83% in *cnep-1Δ,* n = 12; 83% in *cnep-1Δ; lem-2Δ,* n = 6; 0% in *lem-2Δ,* n = 4). These membrane stacks were frequently located near large NE gaps that were only present in the *cnep-1Δ;lem-2Δ* double mutants (3/6 nuclei; Fig. 2 E and Fig. S2 D). The presence of stacked membranes correlated with internal membrane extensions that invaded the nuclear interior (Fig. 2 E, arrows, and Fig. S2 C-E). These data suggest membrane stacks observed by tomograms of nuclear closure may be derived from the ER.

The fact that openings in the NE were observed in *lem-2Δ* mutants deleted for *cnep-1,* suggested that CNEP-1 may be required for the incorporation of ER membranes into NE openings. To test this idea, we imaged an ER marker (SP12::GFP) during post-meiotic NE closure in one-cell embryos. In control embryos, 35% of oocyte-derived pronuclei had fluorescence signal of SP12::GFP that extended from a single region on the nuclear rim into the nucleus (hereafter “internal nuclear membranes”) compared to 0% of sperm-derived pronuclei (Fig. 3 A, left panel and graphs), suggesting that internal nuclear membranes likely arise as the oocyte-derived pronuclei form after meiosis. In the majority of *cnep-1Δ* oocyte-derived pronuclei containing an internal nuclear membrane, the internal nuclear membrane nearly or completely bisected the nucleus (“major” in Fig. 3 A, right panel and graph, and Video 6). *cnep-1Δ* embryos also contained ectopic ER sheets (Fig. 3 A and Video 6, ER clusters in 90%, n = 30 of *cnep-1Δ* nuclei versus 0%, n = 27 of control nuclei), as previously described (Bahmanyar et al., 2014). Thus, the absence of CNEP-1 allows ER membrane sheets to incorporate into the nuclear interior during post-meiotic NE closure.

**Figure 3.**
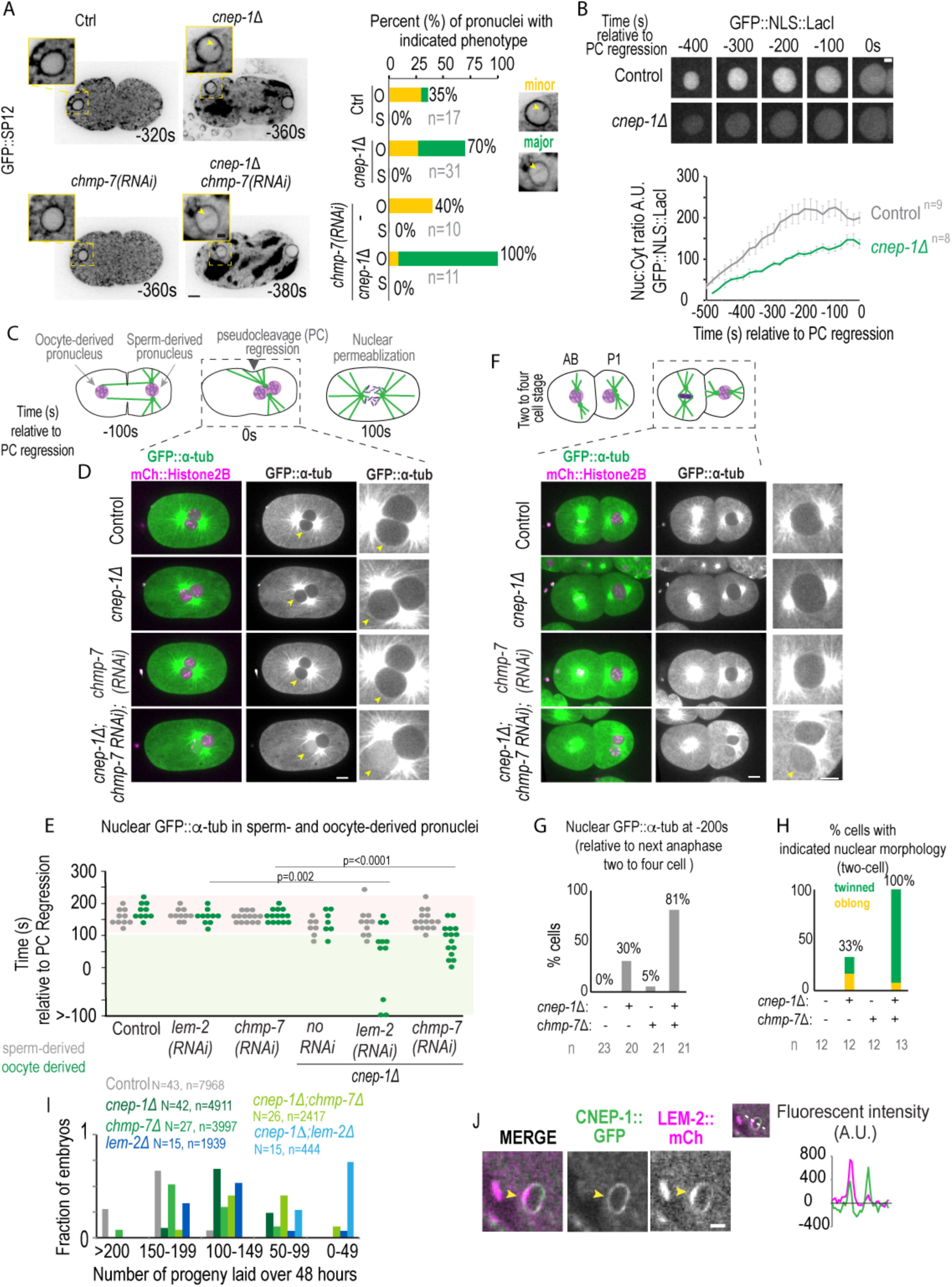
CNEP-1 promotes nuclear closure after meiosis and mitosis independently of LEM-2 and CHMP-7. **(A)** Confocal and magnified images of oocyte-derived pronuclei from time lapse series of oocytes expressing SP12::GFP for the indicated conditions. Plot (right) of percent of oocyte-derived (O) and sperm-derived (S) pronuclei with minor (yellow) or major (green) internal nuclear membranes, n = number of embryos. Scale bars, 5 μm and 2.5 μm. **(B)** Confocal images of oocyte-derived pronuclei from embryos expressing GFP::NLS-LacI in control and *cnep-1Δ* mutants. Scale Bar, 2.5 μm. Below, plot of average +/- SEM of nuclear:cytoplasmic ratio of GFP::NLS-LacI for indicated conditions. Measurements were corrected for background and normalized to nuclear area. n = number of embryos. **(C)** A schematic of pronuclear positioning and NE breakdown in the *C. elegans* zygote. **(D)** Confocal and magnified images from time lapse series of embryos expressing GFP::α-tubulin and mCherry::Histone2B for the indicated conditions. Arrowhead marks the oocyte-derived pronucleus. Scale bar, 5 μm. Time (seconds) is relative to pseudocleavage regression. Scale bars, 5 μm. **(E)** Time (seconds) of entry of GFP::α-tubulin in the sperm-derived (grey) and oocyte-derived pronucleus (green) relative to pseudocleavage regression. **(F)** Confocal images of two-cell embryos at 240 seconds prior to the following anaphase of the P cell (right cell) for the indicated conditions and markers (Scale bar, 10 µm) and magnified images (right) of the P-cell nuclei. Scale bars, 5 μm. **(G)** Plot of percentage of two cell-stage cells with GFP::α-tubulin entry prior to 200 seconds before the following anaphase. N = number of cells **(H)** Plot of percentage of embryos with twinned nuclei or oblong nuclei in the two cell stage. **(I)** Plot of fraction of worms with indicated brood sizes for the indicated conditions over 48 hours, n = embryos, N = adult worms. **(J)** Confocal image of CNEP-1::GFP and LEM-2::mCherry in oocyte pronucleus from time lapse series. Right, plot of background-corrected fluorescent intensities along a line scan.

Consistent with the increase in severity of membrane stacks and extensions in the TEM of thin sections of *cnep-1Δ;lem-2Δ* double mutants (Fig. 2 E), the frequency of major internal nuclear membranes was exacerbated in oocyte-derived pronuclei of *cnep-1Δ* embryos RNAi-depleted of *chmp-7* (Fig. 3 A, bottom right panel and graph). These data suggest that LEM-2 and CHMP-7 limit the invasion of ER membranes that are in excess in *cnep-1Δ* embryos into NE holes during nuclear formation.

Consistent with the idea that internal nuclear membranes in *cnep-1Δ* embryos stem from NE openings facing the meiotic spindle, internal nuclear membranes originate from the GFP::LEM-2 plaque and form when the plaque accumulates in meiosis II (Fig. S2 F and G). Internal nuclear membranes also originate at GFP::LEM-2 plaques of ruptured pronuclei depleted of lamin, and are more severe in ruptured nuclei of embryos also deleted of CNEP-1 (Fig. S2 H). Importantly, oocyte-derived pronuclei in *cnep-1* mutants are unable to retain GFP::NLS-LacI indicating that these nuclei contain openings through which actively imported proteins passively diffuse (Fig. 3 B). Thus, CNEP-1 may limit the production of ER sheets proximal to the NE to control membrane incorporation at NE openings in order to narrow gaps sufficiently to restrict nuclear transport to NPCs.

The enhanced severity of membrane extensions (Fig. 2 E and Fig. 3 A) and the presence of NE gaps (Fig. 2 E) upon deletion of *lem-2* or *chmp-7* in *cnep-1Δ* mutant embryos prompted us to test if these nuclei undergo a relatively greater extent of nuclear-cytoplasmic mixing compared to *cnep-1Δ* mutants alone (Fig. 3 B). To measure an increased severity in nuclear leakiness, we monitored the ability of pronuclei to exclude GFP::α-tubulin, which forms a ∼125 kDa heterodimer with β-tubulin and is actively exported from the nucleus. Soluble GFP::α-tubulin is excluded from control pronuclei during pronuclear migration and rotation and enters the nuclear interior only upon mitotic NE permeabilization (time 120 s relative to psuedocleavage regression, Fig. 3 C-E and Video 7). While most pronuclei in *cnep-1Δ* and RNAi-depleted *chmp-7 or lem-2* embryos exclude GFP::α-tubulin until similar time points as control, nuclear GFP::α-tubulin was observed at significantly earlier time points, and exclusively in the oocyte-derived pronucleus, in *cnep-1Δ* embryos also RNAi-depleted for *chmp-7* or *lem-2* (Fig. 3 D and E, bottom panel and magnified image, and Video 7). Depletion of the ESCRT-III spiral filament protein VPS-32 (Snf7/CHMP4B) did not allow nuclear entry of GFP::α-tubulin in *cnep-1Δ* mutants (Fig. S2 I-K). These data indicate that CNEP-1 and LEM-2/CHMP-7 functions partially overlap to restrict of free diffusion of large proteins across the NE, suggesting that both pathways contribute to NE closure. Furthermore, LEM-2 and CHMP-7 have functions in NE closure that are independent of VPS-32, consistent with reports showing the ability of human LEM2/CHMP7 to co-polymerize *in vitro* (Appen et al., 2019).

The fact that these phenotypes are specific to the oocyte-derived pronucleus indicates that the meiotic spindle increases the susceptibility of the NE to defective remodeling. We predicted that nuclear reformation in the presence of a mitotic spindle would similarly require CNEP-1 and CHMP7 to maintain the NE barrier. GFP::α-tubulin prematurely accessed the interior of both daughter nuclei after mitosis in *cnep-1Δ* embryos RNAi-depleted for *chmp-7* (Fig. 3 F and G). The frequency of twinned nuclei observed at this stage, a phenotype that we previously reported to be caused by excessive ER-NE contacts that delay NE breakdown in *cnep-1Δ* mutants (Bahmanyar et al., 2014), was also enhanced in *cnep-1Δ;chmp-7Δ* double mutants and upon RNAi-depletion of *chmp-7* in *cnep-1Δ* embryos (Fig. 3 F and H). We further corroborated the enhancement of the *cnep-1Δ* phenotypes caused by loss of *chmp-7* by showing that the double mutant has a greater reduction in brood size and an increased percentage of lethal embryos than either single mutant alone (Fig. 3 I and Fig. S2 L). This enhancement in brood size and embryonic lethality was more severe in the *cnep-1Δ;lem-2Δ* double mutant (Fig. 3 I and Fig. S2 L), suggesting that roles for LEM-2 that are independent of CHMP7 support NE remodeling in the absence of CNEP-1.

Thus, ESCRTs protect against passive diffusion of large proteins across the NE in the *cnep-1Δ* background - loss of both pathways has a synergistic effect on the disruption of the NE permeability barrier and the excess internal nuclear membranes – suggesting that ESCRT-III adaptors and CNEP-1 have independent functions during NE closure to prevent major breaches in the NE barrier. Furthermore, our data show that the severe nuclear leakiness and abnormal nuclear morphology caused by loss of both pathways has adverse consequences on germline development and embryogenesis.

Our genetic evidence suggests that the function of CNEP-1 in NE closure is independent of LEM-2 or CHMP-7. Consistent with this finding, CNEP-1 does not accumulate at the GFP::LEM-2 plaque and instead is uniformly localized to the nuclear rim during meiotic NE formation (Fig. 3 J).

The localization of CNEP-1 to the nuclear rim during NE closure in meiosis suggests that it does not function solely at the sites of the forming NE that face the meiotic spindle. Previous work showed that CNEP-1 limits formation of ER sheets near the NE by activating the phosphatidic acid phosphatase, lipin, to bias flux away from PI synthesis (Fig. 4A) (Bahmanyar et al., 2014). Partial RNAi-depletion of lipin caused major internal membrane extensions that bisected the oocyte-derived pronucleus (Fig. S3 A), similar to deletion of CNEP-1. Furthermore, GFP::α-tubulin entered some of the oocyte-derived pronuclei, but not the sperm-derived pronuclei, in embryos RNAi-depleted for lipin in the *chmp-7Δ* mutant (Fig. S3 B and C). These data suggest that CNEP-1’s role in sealing the NE is through activation of lipin.

**Figure 4.**
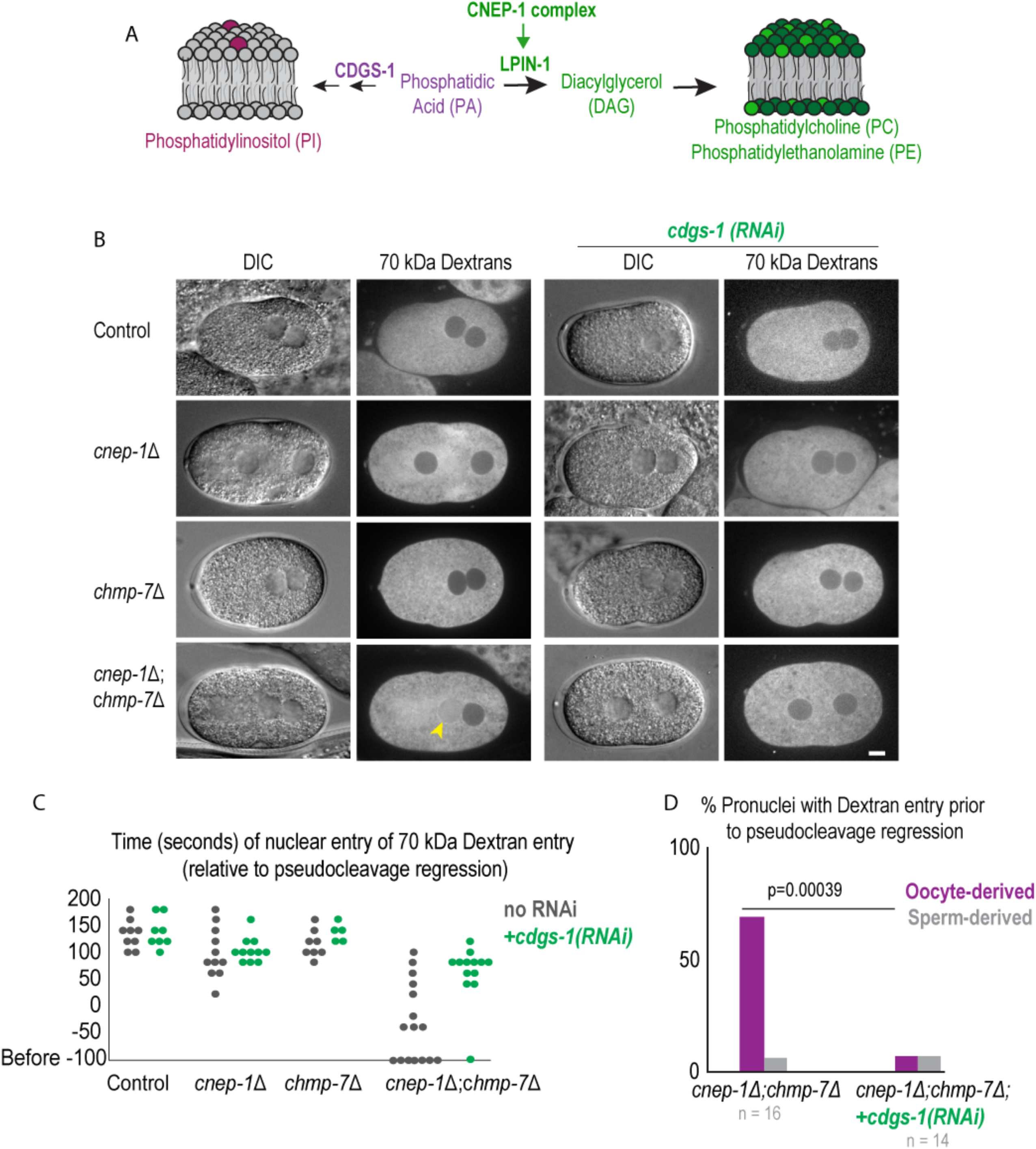
Local control of glycerophospholipid flux by CNEP-1 limit ER sheet production to promote nuclear closure. **(A)** Schematic showing enzymatic pathways to generate phosphatidylcholine/phosphatidylethanolamine or phosphatidylinositol from phosphatidic acid. **(B)** DIC and fluorescence images of embryos from adult worms injected with 70 kDa Texas Red Dextrans. **(C)** Plot of individual times when 70 kDa dextrans enter the oocyte-derived pronucleus relative to pseudocleavage regression for the indicated conditions. **(D)** Percent of oocyte-derived (purple) and sperm-derived (grey) pronuclei with dextran entry prior to pseudocleavage regression for the indicated conditions. A chi-squared test was performed and p-value is reported.

To test if the ectopic ER sheets caused by increased PI levels in *cnep-1Δ* mutants are responsible for the observed defects in the NE permeability barrier, we RNAi-depleted *cdgs-1,* an enzyme in the pathway of PI production that when depleted reduces elevated PI levels and rescues the formation of ectopic ER sheets in *cnep-1Δ* mutants (Fig. 4 A and Fig. S3 D) (Bahmanyar et al., 2014). RNAi-depletion of *cdgs-1* in *cnep-1Δ* and *cnep-1Δ;chmp-7Δ* double mutants rescued ectopic ER sheet formation, as observed by DIC (Fig. 4 B), prevented the diffusion of 70 kDa dextrans into the nuclei of *cnep-1Δ;chmp-7Δ* double mutants (Fig. 4 B-D) and prevented the slightly premature entry of 70 kDa dextrans into the nuclei of *cnep-1Δ* mutants (Fig. 4 C). These data indicate that shifting flux in the phospholipid synthesis pathway towards PI produces ER sheets that are responsible for defects in NE closure in *cnep-1Δ* mutants. The experiments with fluorescent dextrans confirmed that holes in the NE rather than transport defects cause the defects in permeability of the NE in *cnep-1Δ* and *cnep-1Δ;chmp-7Δ* embryos. We conclude that CNEP-1 locally limits PI production to prevent the unrestricted incorporation of ER membranes into NE openings and in turn to promote NE closure.

The acentriolar spindle in *C. elegans* oocyte meiosis provides the unique opportunity to analyze closure of the NE within a region that coincides with spindle microtubules and to determine how a large opening resolves into a sealed NE. Using this system, we identified a role for regulation of phospholipid flux in controlling ER sheet insertion at NE openings to facilitate NE sealing. We propose a model in which LEM-2 and other ESCRT components accumulate around the NE opening surrounding the meiotic spindle to mediate microtubule remodeling (Fig. 5 A, left panel). As the spindle dissipates, ER membranes infiltrate this region and the large opening transitions into many smaller openings that accumulate LEM-2 and ESCRT-III machinery (Fig. 5 A, right panel). At this point, ER membranes move along persisting microtubules to narrow remaining openings such that ER membranes could be targeted specifically to regions where microtubules intersect the NE (Fig. 5B). Then, incorporation of ER membranes narrows NE holes to ensure holes are constricted before ESCRT-III acts (Fig. 5 B). Our data suggest that ER insertion at NE openings provides a mechanism that is sufficient to prevent detectable leakage of proteins across the NE even in the absence of NE-specific adaptors for ESCRT-III (Fig. 5 C).

**Figure 5.**
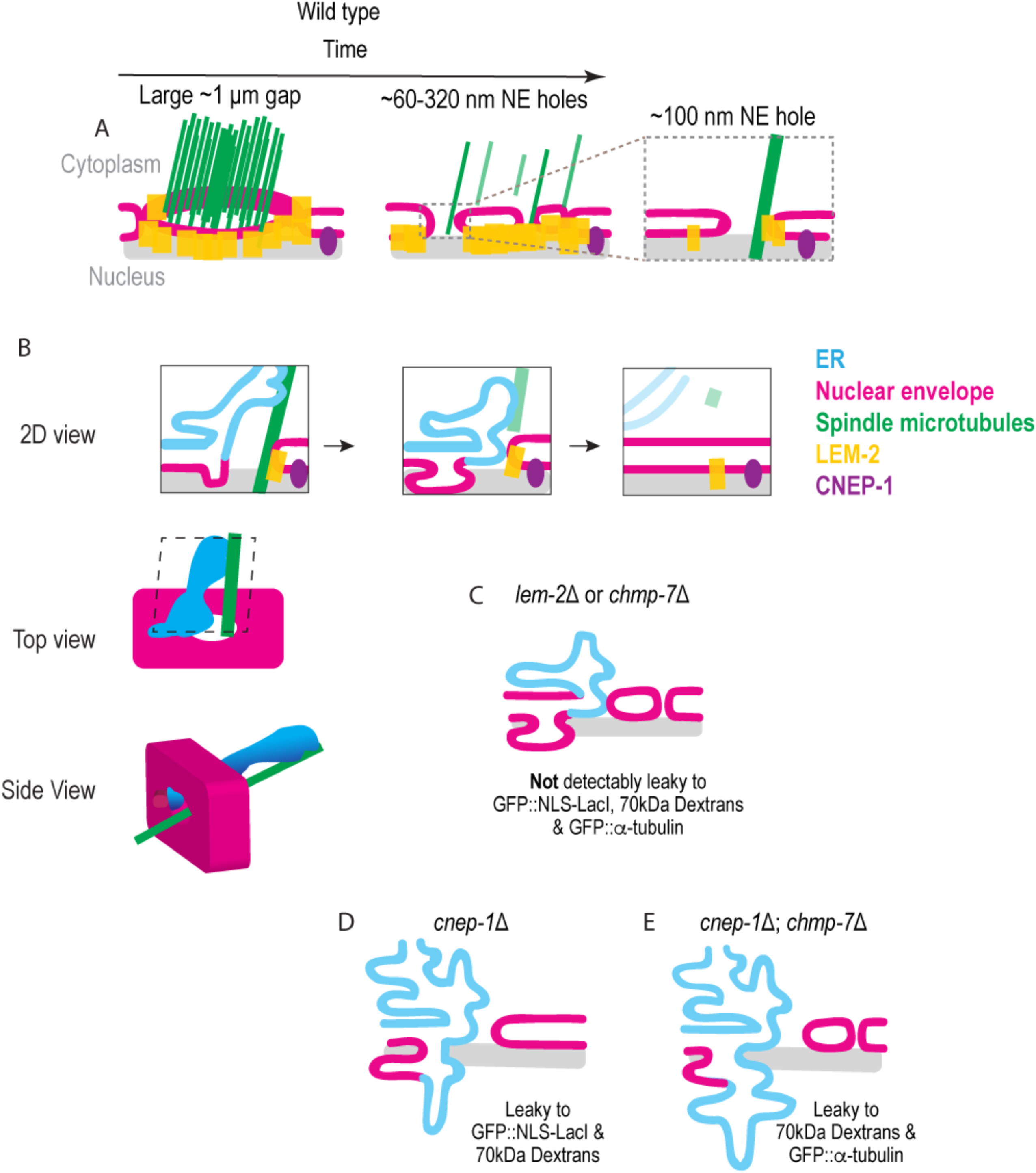
Local regulation of membrane synthesis is coordinated with membrane insertion into nuclear envelope holes to promote nuclear closure. **(A)** LEM-2 accumulates at the rim of the membrane that encloses the meiotic spindle. At a later stage, membranes fill to close the large hole and only a few spindle microtubules persist. **(B)** ER sheets fill in and narrow holes that coincide with spindle microtubules. The membranes of the incoming ER sheet directly feed into and displace the membranes surrounding the hole (left and middle panel). Filling in of holes with incoming membranes of ER sheets and ESCRT-mediated membrane fission seals the NE (right panel). **(C)** In the absence of LEM-2 or CHMP-7, ER membrane sheets fill large holes to restrict the passage of molecules across the membrane. **(D)** In the absence of CNEP-1, the increased production of ER sheets causes an overflow of membranes into NE holes that impedes narrowing of holes. **(E)** Loss of both pathways causes severe leakiness.

The presence of ER sheets at holes in the NE can either help to seal these holes or extend into the nucleus to impair hole closure, such as in the presence of excess ER sheets. Thus, the feeding in of ER sheets to close the NE must be coordinated with the production of ER sheets near the NE. We propose that CNEP-1 enrichment at the NE locally limits the production of proximal ER sheets to prevent the overflow of membranes into NE openings during NE closure (Fig. 5 D). The fact that restoring ectopic ER membrane sheets present in *cnep-1Δ* mutant embryos by shifting flux away from PI production rescues nuclear permeability barrier defects suggests that the defects arise from increased ER sheet production. Thus, it is unlikely that CNEP-1 seals the NE by generation of the fusogenic lipid, DAG (Barona et al., 2005). However, because PI is a precursor to PIPs and PIP2 recruits ESCRT-III associated component, CC2D1B, to the NE (Ventimiglia et al., 2018), it is plausible that higher levels of PI/PIP2 disrupt proper targeting of ESCRT components. Against this idea, our genetic interactions and localization studies indicate that the function of CNEP-1 in NE closure is independent of ESCRT machinery. Thus, our genetic analyses in *cnep-1Δ* mutant embryos together with the observations from electron tomography that ER sheets feed into and narrow NE openings strongly suggest that the increase in ER sheet production caused by elevated levels of PI causes defects in the NE permeability barrier.

Without either NE sealing pathway, the NE permeability barrier is severely disrupted (Fig. 5 E) – the fact that large proteins traverse the NE under this condition suggests either the number or the size of the gaps are affected. Because depletion of VPS-32/CHMP4B did not cause major breaches in the NE, our data suggest that ESCRT components LEM-2 and CHMP-7 have functions independent of recruiting VPS-32. LEM-2 and CHMP-7 may be involved in constricting NE holes to prevent major NE sealing defects caused by membrane overflow in the absence of CNEP-1. The fact that depletion of *chmp-7* in the *cnep-1Δ* background increases the frequency of internal nuclear membranes after meiosis as well as the percentage of twinned nuclei after mitosis also suggests a potential role for ESCRTs in membrane clearance, which has been proposed as a role for ESCRTs in cytokinetic abscission (König et al., 2017).

The idea that redundant parallel membrane sheets nearby NE openings (Fig. 2 D and E) are nuclear membranes displaced by incoming ER sheets is consistent with the observation that annulate lamellae inserted at NE openings displace nuclear membranes during nuclear expansion in *Drosophila* embryos. Interestingly, insertion of annulate lamellae does not disrupt the NE permeability barrier (Hampoelz et al., 2016). Thus, ER sheet insertion into NE openings during NE closure might similarly ensure NE holes are covered, providing a way to restrict passage across NE holes independent of membrane fission. Whether ER sheet insertion is also able to restrict passage through NE holes in other systems to narrow large openings is not known.

Because CNEP-1 also limits membrane incorporation at NE rupture sites, ER membranes likely also contribute to repair of NE holes after rupture. However, given there are no spindle microtubules, it is unclear how these membranes would be delivered specifically to these NE openings. It is plausible that transient changes in NE organization at NE rupture sites may signal or support membrane insertion (De Vos et al., 2011; Denais et al., 2016; Raab et al., 2016; Penfield et al., 2018). For instance, BAF crosslinks chromatin at the core region of chromatin that coincides with spindle microtubules (Samwer et al., 2017) and was recently shown to accumulate at rupture sites (Halfmann et al., 2019), providing a possible mechanism to direct membrane incorporation to this region. Identifying how ER sheets are delivered into NE holes in different contexts is an important next step.

Together our data demonstrate ER sheet insertion as a mechanism that functions independently of ESCRT-membrane remodeling to close NE holes during oocyte meiosis and mitosis. Because the NE and ER are contiguous membrane systems, it has been unclear how each organelle establishes and maintains its unique structure. Our findings emphasize the importance of local regulation of phospholipid flux to form and remodel the NE from ER-derived membranes.

## Acknowledgements

We thank Kim Gibson for advice and training on IMOD to build 3D model of tomography data and the Xinran Liu and Morven Graham (Yale EM Facility) for assistance with 2D transmission electron microscopy of thin section. We are grateful to the members of the Bahmanyar lab for helpful discussion and for Tom Pollard for feedback on the manuscript. This work was supported by an NSF CAREER Award to SB (NSF CAREER 1846010), NSF award #1661900 to AA, and NIH R01GM088151 to AA, and by the German Research Foundation (DFG grants MU1423/3-2 and 4-1 to TMR).

## Author Contributions

L. Penfield and S. Bahmanyar conceived the project. L. Penfield performed the majority of the experiments. E. Szentgyörgy and T. Müller-Reichert prepared samples by high-pressure freezing and generated electron tomograms. L.P. generated 3D models of reconstructed tomograms. L. Penfield and R. Shankar generated strains used in the manuscript. R. Shankar prepared samples for thin-section transmission electron microscopy. A. Laffitte did live imaging of some mitotic embryos and M. Mauro provided results that guided experiments. L. Penfield and S. Bahmanyar wrote the manuscript, with input from all authors. S. Bahmanyar supervised the project.

## Materials and Methods

### Strains

Strains were maintained on nematode growth media plates seeded with OP50 *Escherichia coli* and stored at 20°C. Strains used in this paper are listed in Key Resources Table. The deletion of T24B8.2 (*chmp-7)* was generated with CRISPR-Cas9 with crRNAs listed in the Key Resources Table. To generate the T24B8.2 deletion by CRISPR-Cas9, we injected Cas9-CRISPR RNA (crRNA) *trans*-activating crRNA (tracrRNA) ribonucleoprotein complexes that were assembled *in vitro* into *C. elegans* gonads (Paix et al., 2015). In brief, crRNAs were annealed with trRNAs at 95 C for 2 minutes, and the annealed complexes (final concentration: 11.7 uM) were added to Cas9 protein (qb3 Berkeley, final concentration: 14.7 uM), along with *dpy-10* crRNAs annealed with trRNA (final concentration: 3.7 uM) and a *dpy-10*(cn64) repair template (final concentration: 29 ng/mL) as a co-CRISPR positive control (Arribere et al., 2014). The mixture was also injected into *cnep-1Δ* adult worms to generate double mutants. Roller F1 progeny were singled out onto OP50 plates, allowed to lay progeny, and genotyped by PCR to identify a strain with a deletion for T24B8.2. Deletion strains were outcrossed to N2 worms at least 4X after their generation.

### RNA interference

dsRNAs used in this study were generated with the oligos listed in the Key Resources Table. T3 and/or T7 transcription reactions (MEGAscript, Life Technologies) were ran with purified PCR products as templates for 5 hours. T3 and T7 reactions were mixed and purified using a phenol-chloroform purification for dsRNAs. For depletion of a gene, the dsRNA was diluted to 1 mg/mL in 1X soaking buffer (Penfield et al., 2018) and spun down in a centrifuge for 30 minutes at 4 °C. dsRNA was loaded into an injecting needle and injected into L4 worms. Worms were incubated at 20°C for 24 hours (*chmp-7, lem-2, lmn-1, vps-32)*, 48 hours (*cdgs-1, vps-32)*, or 12 hours (*lpin-1)* prior to imaging.

### Brood Size and Lethality Tests

L4 worms were isolated onto individual OP50-seeded plates and allowed to lay progeny for 24 hours. Then, the worms were moved to individual plates and allowed to lay progeny for an additional 24 hours and removed from the plates. Embryos were allowed one day to hatch prior to counting the number of hatched and unhatched embryos from each plate. The number of embryos were combined over the 48 hours for the brood size and embryonic lethality quantification.

### Live and fixed microscopy

Adult hermaphrodites were dissected in M9 to release embryos for live microscopy. Embryos were transferred to a 2% agarose slide compressed by a cover slip, and imaged at room temperature (∼21°C). Live and fixed embryos were imaged on an inverted Nikon (Melville, NY) Ti microscope equipped with a confocal scanner unit (CSU-XI, Yokogawa) with solid state 100-mW 488-nm and 50-mW 561-nm lasers, with a 60 × 1.4 NA plan Apo objective lens, and a high-resolution ORCA R-3 Digital CCD Camera (Hamamatsu). Early embryos for live imaging were imaged at a temporal resolution of 20 s and five z-slices were acquired with 2-μm steps. For 3D-projections (Fig. 1 D and Fig. S1 E),10-15 z-slices were acquired with 0.5 μm steps.

### Dextrans

Fluorescent Dextrans (Texas Red™ 70,000 MW, Lysine Fixable, Thermo Fischer Scientific) were diluted in 1X soaking buffer to 0.8 mg/mL as previously described (Portier et al., 2007) and injected into gonads of young adults. Injected worms were incubated at 20 °C for 5 hours prior to imaging.

### Immunofluorescence

Immunofluorescence was performed as described previously (Penfield et al., 2018). Primary antibodies in PBST were incubated overnight at 4°C in a humid chamber (45 µl per slide; mouse α-tubulin [clone DM1A; EMD Millipore], 0.5 µg/ml; goat α-GFP [Hyman lab], 1 µg/ml; rabbit α-VPS-32 [Audhya lab], 0.5 µg/ml). Following primary antibody incubation, slides were washed twice for 10 min in PBST and incubated for 1 hour in the dark with secondary antibody in PBST at room temperature (anti-rabbit Cy3/Rhodamine, 1:200; anti-mouse Cy5 1:200; anti-goat FITC 1:200; anti-rabbit Cy5 1:200). All secondary antibodies were purchased from Jackson Immunoresearch. Slides were then washed with PBST twice for 10 min in the dark and mounted with Molecular Probes ProLong Gold Antifade Reagent with DAPI.

### Immunoblot

For each condition, 35 adult worms were collected and washed two times with M9 (Na2HPO4, 4.2 mM; KH2PO4, 2.2 mM; MgSO4, 1 mM; and NaCL, 8.6 mM in ddH2O) and 0.1% Triton and brought to a volume of 30 µl in microcentrifuge tubes. Then, 10 µl of 4X sample buffer were added to each tube and tubes were sonicated for 10 min at 70°C. Then, tubes were incubated at 95°C for 5 min followed by an additional 10 minutes of sonication at 70°C. The samples were frozen at –20°C prior to running a protein gel. 20 µl (∼17.5 worms) of lysate was loaded into each lane and were run on an SDS–PAGE gel. Samples were then transferred to a nitrocellulose membrane for immunoblotting; primary antibodies were added at a concentration of 1 µg/ml for rabbit-anti-VPS-32 (Audhya lab) and mouse-anti-α-tubulin (EMD Millipore). Secondary antibodies were diluted 1:5000 for horseradish peroxidase (HRP)-conjugated goat-anti-rabbit and 1:7000 for HRP conjugated goat-anti-mouse (ThermoFischer Scientific)

### Image analysis

Fluorescence intensity measurements were made using ImageJ (Fiji) software (National Institutes of Health), and measurements were analyzed in a systematic manner to minimize bias. All embryos were included for the analysis unless they became arrested/were damaged.

To measure the nuclear:cytoplasmic ratio of GFP::NLS::LacI, the average nuclear and cytoplasmic signals were measured and subtracted for the average camera background for each time point. The nuclear:cytoplasmic ratio was then normalized for nuclear area for each time point.

For line scans, a five-pixel-wide, 10-μm-long line was drawn across the forming oocyte-derived pronucleus and each fluorescent intensity value was plotted along a line after subtraction of background (the average of a five-pixel-wide, 10-μm-long line in the background). To measure the percent of fluorescent intensity of the meiotic spindle (Fig. 1), 5 pixel wide line scans were drawn across the spindle at time 0 and the time of GFP::LEM-2 plaque formation and subtracted for the average camera background. The average maximum values of the time of GFP::LEM-2 plaque formation were divided by the average maximum values of time of maximum spindle shortening.

### Transmission Electron Microscopy

Whole *C. elegans* animals were mounted on 1-hexadecene coated Type A aluminum sample holders (100 μm deep), in a paste of OP50 bacteria and frozen using a Balzers HPM 010 high pressure freezer. The samples were rapidly freeze substituted (∼4 hours) in 1% OsO4, 1% H2O in acetone (McDonald, 2014). Animals were infiltrated with increasing concentration of epoxy resin (EMbed 812, EMS) and polymerized at 60°C for 24 hours. Samples were then remounted on blank resin blocks for sectioning. Hardened blocked were sectioned using a Leica UltraCut UC7. 60 nm sections were collected on formvar coated nickel grids and stained using 2% uranyl acetate and lead citrate. 60nm Grids were viewed FEI Tencai Biotwin TEM at 80Kv. Images were taken using Morada CCD and iTEM (Olympus) software.

### Electron Tomography and 3D modeling

Hermaphrodites were dissected in Minimal Edgar’s Growth Medium (Edgar, 1995; Woog et al., 2012) and embryos in meiosis were transferred to cellulose capillary tubes (Leica Microsystems, Vienna, Austria). The embryos were observed with a stereo microscope and placed into membrane carriers at anaphase I for immediate cryo-immobilization using an EMPACT2+RTS high-pressure freezer (Leica Microsystems, Vienna, Austria) (Redemann et al., 2018). Freeze substitution was performed over 3 d at −90°C in anhydrous acetone containing 1% OsO4 and 0.1% uranyl acetate using an automatic freeze substitution machine (EM AFS, Leica Microsystems, Vienna, Austria). Epon/Araldite infiltrated samples were embedded in a thin layer of resin and polymerized for 3 d at 60°C. Serial semi-thick (300 nm) sections were cut using an Ultracut UCT Microtome (Leica Microsystems, Vienna, Austria), collected on Formvar-coated copper slot grids and post-stained with 2% uranyl acetate in 70% methanol followed by Reynold’s lead citrate (Müller-Reichert et al., 2007). For dual-axis electron tomography (Mastronarde, 1997), 15-nm colloidal gold particles (Sigma-Aldrich) were attached to both sides of the semi-thick sections. Series of tilted views were recorded using a TECNAI F30 transmission electron microscope (Thermo Fisher Scientific) operated at 300 kV. Images were captured every 1.0° over a ±60° range at a pixel size of 2.3 nm using a Gatan US1000 2K x 2K CCD camera. Using the IMOD software package (Kremer et al., 1996) (http://bio3d.colorado.edu/imod), montages of 2 x 1 were collected and combined for each serial section (Redemann et al., 2018). Tomograms were computed for each tilt axis using the R-weighted back-projection algorithm (Gilbert, 1972). The IMOD software package was also used to segment microtubules, the ER and nuclear membranes (Kremer et al., 1996).

### Statistical Analysis

All data points are reported in graphs and data analysis, middle and error bar types are noted in figure legends. Statistical analysis was performed on data sets with multiple samples and independent biological repeats. A nonparametric Mann Whitney U test was used to compare data sets without a normal distribution, determined by a Shapiro-Wilk test, while a chi-squared test was utilized to compare fraction of embryos with indicated phenotypes between conditions. The type of test utilized, sample sizes and p-values are reported in figure legends or in text (p<0.05 defined as significant).

