## Supplemental Material for "Local regulation of lipid synthesis controls ER sheet insertion into nuclear envelope holes to complete nuclear closure"

**Inventory of Supplemental Materials**

Figures S1, S2, S3

Videos 1-7

Supplemental Figure Legends

Video Legends

Key Resources Table

#### Supplemental Figure Legends

**Figure S1. Nuclear membranes asymmetrically enclose oocyte chromatin during nuclear formation after meiosis II,** related to Figures 1 and 2. **(A)** Table of time in seconds (average +/- SD) of indicated events relative to chromosome segregation (anaphase II onset) or maximal spindle shortening. n = number of embryos. **(B)** Time in seconds of the duration of appearance to dispersal of GFP::LEM-2 plaque. Average +/- SD is shown. **(C)** Fold enrichment of GFP::LEM-2 compared to opposite side of the nuclear envelope at 240 seconds after anaphase onset. Average +/- SD is shown. **(D)** Confocal images from a time lapse series of *C. elegans* oocyte meiosis II expressing SP12::GFP and mCherry::Histone2B. Representative example of n = 5 embryos is shown. Time is in seconds relative to anaphase II onset. **(E)** Single z-slice, max z-projection, and a 3D-projection of rupture site in a sperm-derived pronucleus depleted of lamin expressing GFP::LEM-2. Arrowheads mark a gap between the GFP::LEM-2 plaque that has formed at a NE rupture site. **(F)** Fixed *C. elegans* embryo stained for GFP::LEM-2 (magenta) and α-tubulin (green). Magnified images on the left are of the oocyte-derived pronucleus. **(G)** CRISPR strategy used to generate deletion in T24B8.2 *(chmp-7)* locus, a gene predicted to encode the ortholog of human CHMP7. crRNAs used are shown in green. **(H)** Tomographic slice of *C. elegans* oocyte in the early stage of anaphase. Scale bar, 500 nm. End on views (top) and side view (bottom) of 3D model of the NE (magenta) rendered from tracing of NE in tomographic slices in which entire gap is shown. Scale bar, 250 nm. Yellow arrowheads mark flat, membrane sheets that are continuous with region of the NE that partially fills the gap.

**Figure S2. CNEP-1 restricts nuclear membrane incorporation during NE sealing processes and genetically interacts with NE adaptors for ESCRT-III,** related to Figures 2 and 3. **(A-B)** Schematics of the *cnep-1* (A) or *lem-2* (B) gene loci and the deletion alleles utilized in this study. **(C)** Transmission electron micrographs of thin sections of nuclei for the indicated conditions and images (below) where the NE (magenta) and ER membranes (blue) are traced.

Scale bar, 1 μm. Magnified examples for first three images are also shown in Fig. 2 E. Yellow arrowheads mark membrane stacks, green arrow marks internal membrane extension, and red asterisks mark gaps in the NE. **(D)** Magnified TEM of thin section of nucleus in (C) of *lem-2Δ; cnep-1Δ* mutant embryos. Scale bar, 500 nm. The NE (magenta) is traced (right). **(E)** Plot of percent of cells with nuclei with internal nuclear membranes scored from TEM thin sections for the indicated conditions. n = number of cells. **(F)** Confocal images from a time lapse series of oocytes expressing GFP::LEM-2. Green arrowhead marks GFP::LEM-2 plaque and yellow arrowheads mark internal membrane extension. Scale bar, 2.5 μm. **(G)** Plot of the percent of nuclei with GFP::LEM-2 marked internal nuclear membranes at the time of NE formation after meiosis II. **(H)** Confocal images of ruptured oocyte-derived (left) and sperm-derived (right) pronuclei from time lapse series of embryos expressing GFP::LEM-2 and percentages (right) of pronuclei that have internal nuclear membranes marked with GFP::LEM-2 for the indicated conditions. Yellow arrowheads mark internal membrane extensions. n = number of embryos. **(I)** Immunoblot probing for tubulin and VPS-32 for the indicated conditions. **(J)** Confocal images for the indicated conditions of pronuclei 80 seconds after pseudocleavage cleavage regression. Arrowhead marks oocyte-derived pronucleus. **(K)** Time (seconds) of entry of GFP::α-tubulin in the sperm-derived (grey) and oocyte-derived pronucleus (green) relative to pseudocleavage regression for the indicated conditions. **(L)** Plot of the fraction of adult worms with the indicated % embryonic lethality ranges over 48 hours. N = number of adult worms. n = number of embryos.

**Figure S3. Depletion of lipin causes internal nuclear membrane and loss of the nuclear permeability barrier in the oocyte-derived pronucleus**, related to Figures 3 and 4. **(A)** Confocal images of *C. elegans* embryos -180 seconds before pseudocleavage regression (PC regression) expressing SP12::GFP (an ER marker) from a time lapse series. Scale bar, 5 μm. Magnified images (right) are of oocyte-derived pronuclei. Arrowhead marks internal membrane extension that bisects the nucleus. Scale bar, 2.5 μm. Plot (right) of percentage of embryos with "major" and "minor" internal nuclear membranes for indicated conditions. **(B)** Confocal images from time lapse series of control and lipin RNAi-depleted *C. elegans* embryos expressing GFP::α-tubulin at pseudocleavage regression. Arrowhead marks oocyte-derived pronucleus with nuclear soluble GFP::α-tubulin, Scale bar, 5 μm. **(C)** Plot of times in seconds of the onset of entry of soluble GFP::α-tubulin into the oocyte-derived pronucleus. Average +/- SD is shown. Time in seconds is relative to when soluble GFP::α-tubulin enters the sperm-derived pronucleus. **(D)** Representative DIC and fluorescence images of embryos expressing SP12::GFP for the indicated conditions.

**Supplemental Video Legends**

**Video 1. LEM-2 plaque forms and disperses as the spindle dissipates and detaches from the oocyte-derved pronucleus, related to Figure 1.** *C. elegans* oocyte in meiosis II expressing GFP::LEM-2 and mCherry::α-tubulin. Arrows mark initial GFP::LEM-2 rim and the formation of the GFP::LEM-2 plaque. Movie starts at -100s prior to maximal spindle shortening and mages were acquired every 20 seconds and the playback rate is 60X real time. Scale bar, 10 μm

**Video 2. Electron tomogram of oocyte-derived pronucleus in mid-anaphase, related to Figure 2.** Region of electron tomogram of *C. elegans* oocyte in mid-anaphase that was traced in Figure 2A. Playback rate is 20 z-slices per second. Scale bar, 500 nm.

**Video 3. 3D model of region of tomogram in Movie S2, related to Figure 2.** The NE (magenta) is initially shown from the side farthest from the microtubules (green) and is rotated 90 degrees to a birds eye view where the top is the region of the NE facing the meiotic spindle.

The model is then magnified to show the large gap in the NE that intersects the meiotic spindle. Scale bar in all frames, 250 nm.

**Video 4. Electron tomogram of oocyte-derived pronucleus in late-anaphase, related to Figure 2.** Region of electron tomogram of *C. elegans* in late anaphase that is shown and traced in Figure 2B. Playback rate is 10 z-slices per second. Scale bar, 100 nm.

**Video 5. 3D model of region of tomogram in Movie S4.** The NE (magenta) is oriented with the region facing the extruding polar body at the top of the image. The model is rotated 90 degrees and magnified on a NE hole occluded by a remaining microtubule (green). A membrane sheet (blue) contacting the outer nuclear membrane feeds into and narrows the hole in the NE. Scale bar in all frames, 250 nm.

**Video 6. Internal nuclear membranes after oocyte meiosis II in *cnep-1Δ* embryos.** A control *C. elegans* embryo (left) and a *cnep-1Δ* embryo (right) expressing SP12::GFP imaged from shortly after meiosis. Arrow marks internal nuclear membrane. Images were acquired every 20 seconds and the playback rate is 60X real time, Scale bar, 10 μm

**Video 7. CNEP-1 and CHMP7 maintain the NE permeability barrier after oocyte meiosis.** *C. elegans* embryos expressing GFP::α-tubulin for the indicated conditions imaged shortly after meiosis up until NE breakdown. Arrow marks GFP::α-tubulin entry into oocyte-derived pronucleus in *cnep-1Δ;chmp-7(RNAi)* embryos. Images were acquired every 20 seconds and the playback rate is 60X real time, Scale bar, 10 μm

**Key Resources Table**

**Strains used in this study**

| **Strain Name** | **Strain genotype** |
|---|---|
| N2 (Bristol) | Wild-type (ancestral) |
| OD83 | unc-119(ed3) III; ltIs37 [pAA64; Ppie-1/mCherry::his-58; unc119(+)] IV. qaIs3507 [pie-1::GFP::lem-2; unc-119(+)] |
| OD1344 | scpl-2(tm4369)II; unc-119(ed3) III; ltIs37 [pAA64; Ppie-1/mCherry::his-58; unc119(+)] IV. qaIs3507 [pie-1::GFP::lem-2; unc-119(+)] |
| SBW71 | qaIs3507 [pie-1::GFP::LEM-2 + unc-119(+)]; weIs21 [pJA138 (pie-1::mCherry::tub::pie-1)] |
| SBW47 | unc-119(ed3) III; ltIs37 [pAA64; pie-1/mCHERRY::his-58; unc-119 (+)] IV;ltIs75 [(pSK5) pie-1::GFP::TEV-Stag::LacI + unc-119(+)]. |
| SBW65 | scpl-2(tm4369)II;unc-119(ed3) III; ltIs37 [pAA64; pie-1/mCHERRY::his-58; unc-119 (+)] IV;ltIs75 [(pSK5) pie-1::GFP::TEV-Stag::LacI + unc-119(+)]. |
| MSN772 | *chmp-7*(hz12) II |
| SBW79 | *chmp-7*(hz12) II ; unc-119(ed3) III; ltIs37 [pAA64; pie-1/mCHERRY::his-58; unc-119 (+)] IV;ltIs75 [(pSK5) pie-1::GFP::TEV-Stag::LacI + unc-119(+)]. |
| MSN224 | lem-2(tm1582) II |
| SBW59 | lem-2(tm1582) II;scpl-2(tm4369)II |
| OD417 | scpl-2(tm4369)II 4X outcross |
| SBW54 | bqSi226 [lem-2p::lem-2::mCherry + unc-119(+)] IV; unc-119(ed3) III; ltSi185 [pSB67; scpl-2::scpl-2::GFP; unc-119 (+)]I |
| SBW32 | ltIs24 [(pAZ132) pie-1p::GFP::tba-2 + unc-119(+)];unc-119(ed3) III; ltIs37 [pAA64; pie-1/mCHERRY::his-58; unc-119 (+)] IV |
| SBW49 | ltIs24 [pAZ132; pie-1/GFP::tba-2; unc-119 (+)]; unc-119(ed3) III; ltIs37 [pAA64; pie-1/mCHERRY::his-58; unc-119 (+)] IV; scpl-2(tm4369)II |
| SBW63 | chmp-7(hz12) II; ltIs37 [pAA64; pie-1/mCHERRY::his-58; unc-119 (+)] IV; unc-119(ed3) III; ltIs24 [pAZ132; pie-1/GFP::tba-2; unc-119 (+)] |

| | |
|---|---|
| | (SBW58XSBW32) |
| SBW73 | chmp-7(sbw2)II;cnep-1(tm4369)II 4X outcross |
| SBW75 | *chmp-7* (sbw2) II; cnep-1(tm4369)II; ltIs37 [pAA64; pie-1/mCHERRY::his-58; unc-119 (+)] IV; unc-119(ed3) III; ltIs24 [pAZ132; pie-1/GFP::tba-2; unc-119 (+)] |
| OD270 | unc-119(ed3) III; ojls23[SP12::GFP unc-119(+)]; ltIs37 [pAA64; pie-1/mCHERRY::his-58; unc-119 (+)] |
| SBW66 | scpl-2(tm4369)II; unc-119(ed3) III; ojls23[SP12::GFP unc-119(+)]; ltIs37 [pAA64; pie-1/mCHERRY::his-58; unc-119 (+)] |

**CRISPR crRNAs**

| **Target** | **crRNA sequence** |
|---|---|
| T24B8.2_ start | CUUCCUAUCUCCCUUCCGAA |
| T24B8.2_end | CAAAGAAGUAACGCCAGAAG |

**Oligos**

| **Name** | **Sequence** |
|---|---|
| dsRNA T3 forward *lem-2* | AATTAACCCTCACTAAAGGAGAAAATGTCGGATGCAGAG |
| dsRNA T7 forward *lem-2* | TAATACGACTCACTATAGGTTGTTAGGCGTCGAAGAAAC |
| dsRNA T7 forward T24B8.2(*chmp-7*) | TAATACGACTCACTATAGGTCGGTGAATGGAGAGATCGT |
| dsRNA T7 reverse T24B8.2 (*chmp-7*) | TAATACGACTCACTATAGGGTTCTGAGCACGTCCTTTGT |
| dsRNA T3 *lpin-1* | AATTAACCCTCACTAAAGGGGCCATTTGTCCACTCTCAT |
| dsRNA T7 *lpin-1* | TAATACGACTCACTATAGGCTTACACACTCGGCGGTTTT |

| | |
|---|---|
| dsRNA T7 forward *cdgs-1* | TAATACGACTCACTATAGGTGTTTGGATTCTTTTGGGGA |
| dsRNA T7 reverse *cdgs-1* | TAATACGACTCACTATAGGTTTGAAGCATCAGGAACACG |
| dsRNA T3 forward *lmn-1* | AATTAACCCTCACTAAAGGCCTCGATTCGGCTCAAGAT |
| dsRNA reverse T7 *lmn-1* | TAATACGACTCACTATAGGTTCTCCACCACCAATTTATCG |
| dsRNA forward T7 *vps-32* | TAATACGACTCACTATAGGCAGTTGGCCCATATTGACGG |
| dsRNA reverse T7 *vps-32* | TAATACGACTCACTATAGGGAGTATCCGGAAGCGTGACT |

**Penfield et al. Figure S1**

A

Average time of event +/- SD (seconds)

| | Initial LEM-2 NE accumulation | LEM-2 punctum forms | LEM-2 punctum resolves | Spindle detaches from pronucleus |
|---|---|---|---|---|
| relative to anaphase II onset | **28 +/- 14s** n=9 | **182 +/- 32s** n=9 | **388+/- 74s** n=9 | **n/a** n=9 |
| relative to maximum spindle shortening | **2 +/- 11s** n=9 | **147 +/- 30s** n=10 | **353 +/- 76s** n=9 | **305 +/- 69s** n=8 |

B

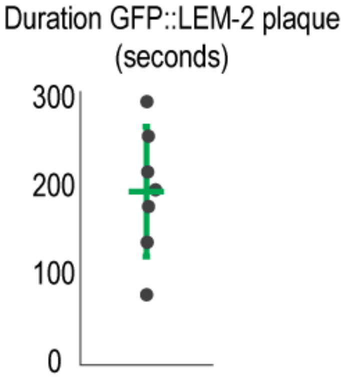


C

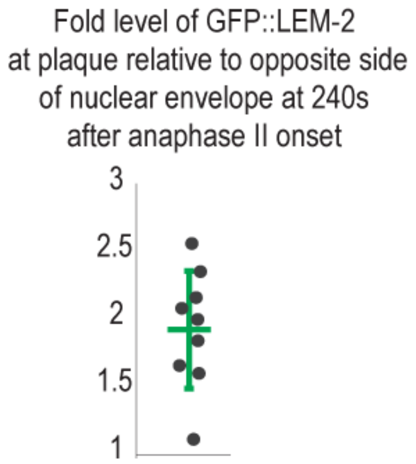


D

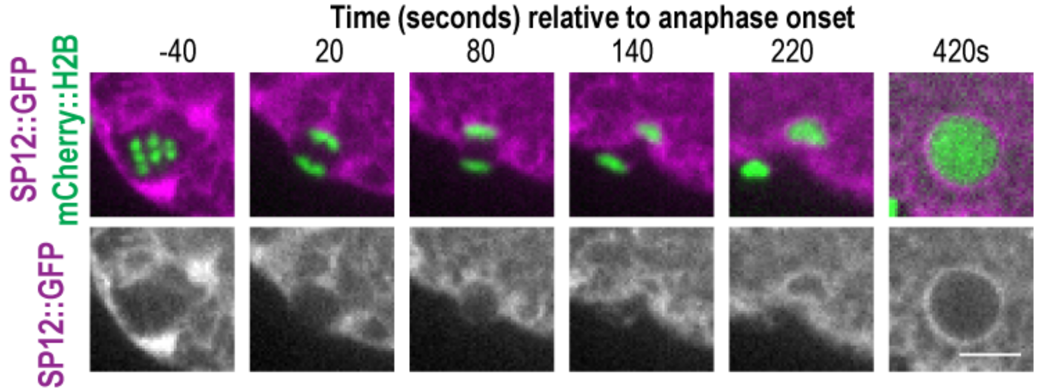


E

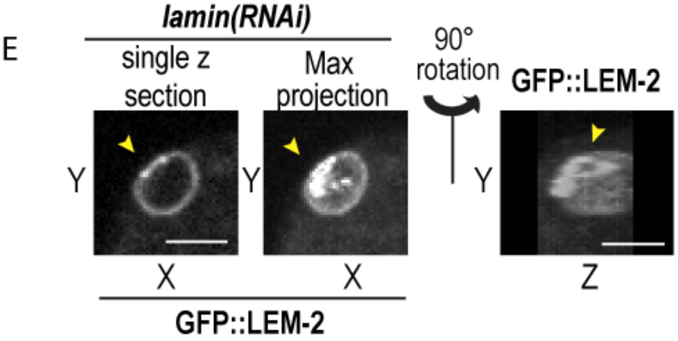


F

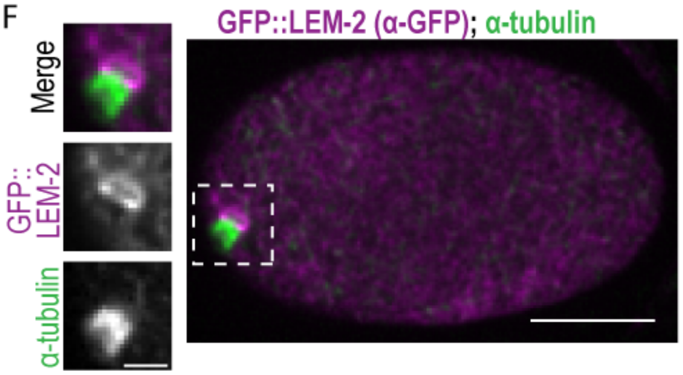


G

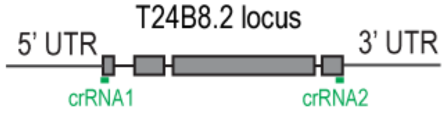


H

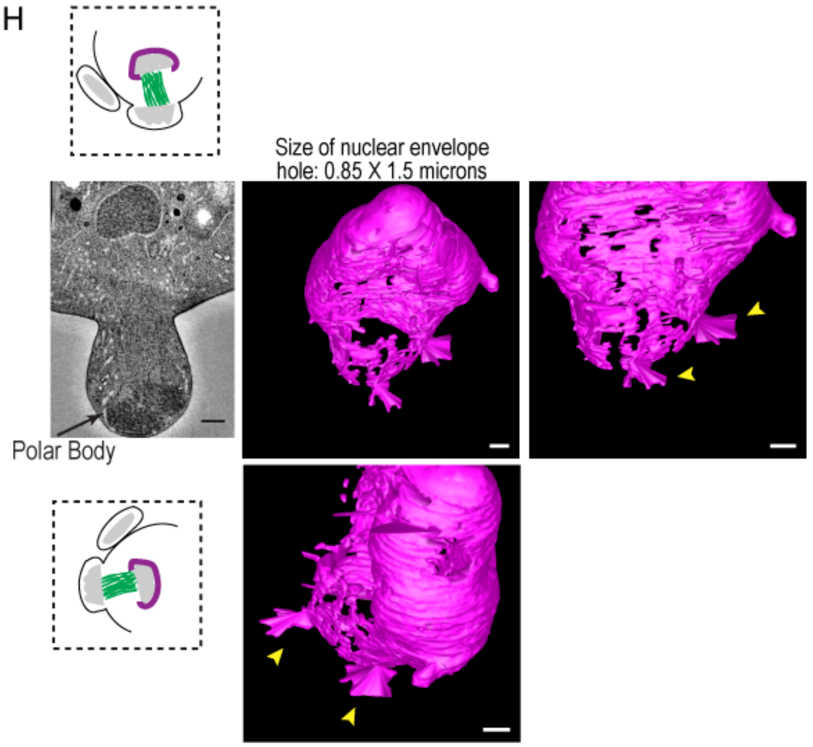

Penfield et al. Figure S2

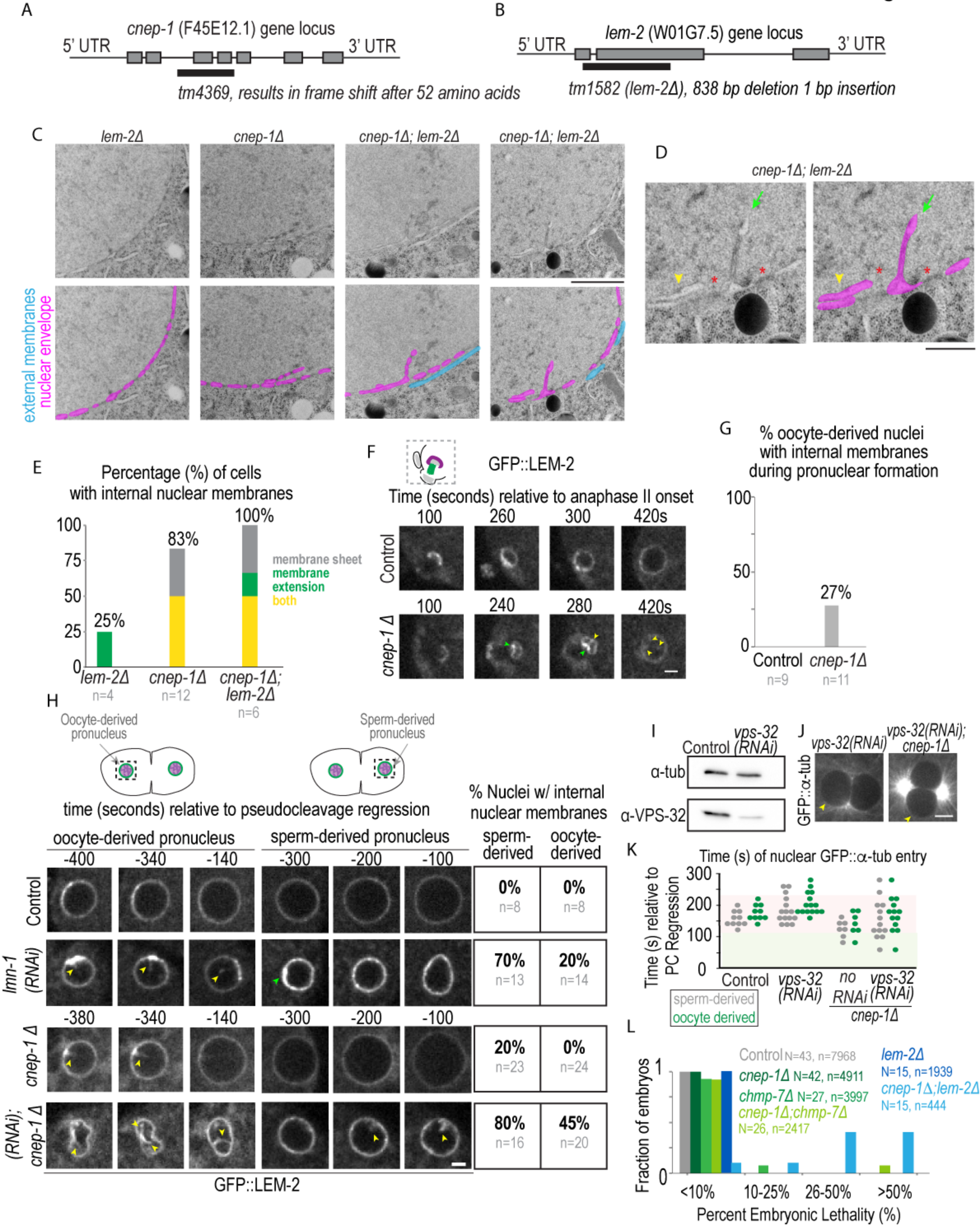

### Penfield et al. Figure S3

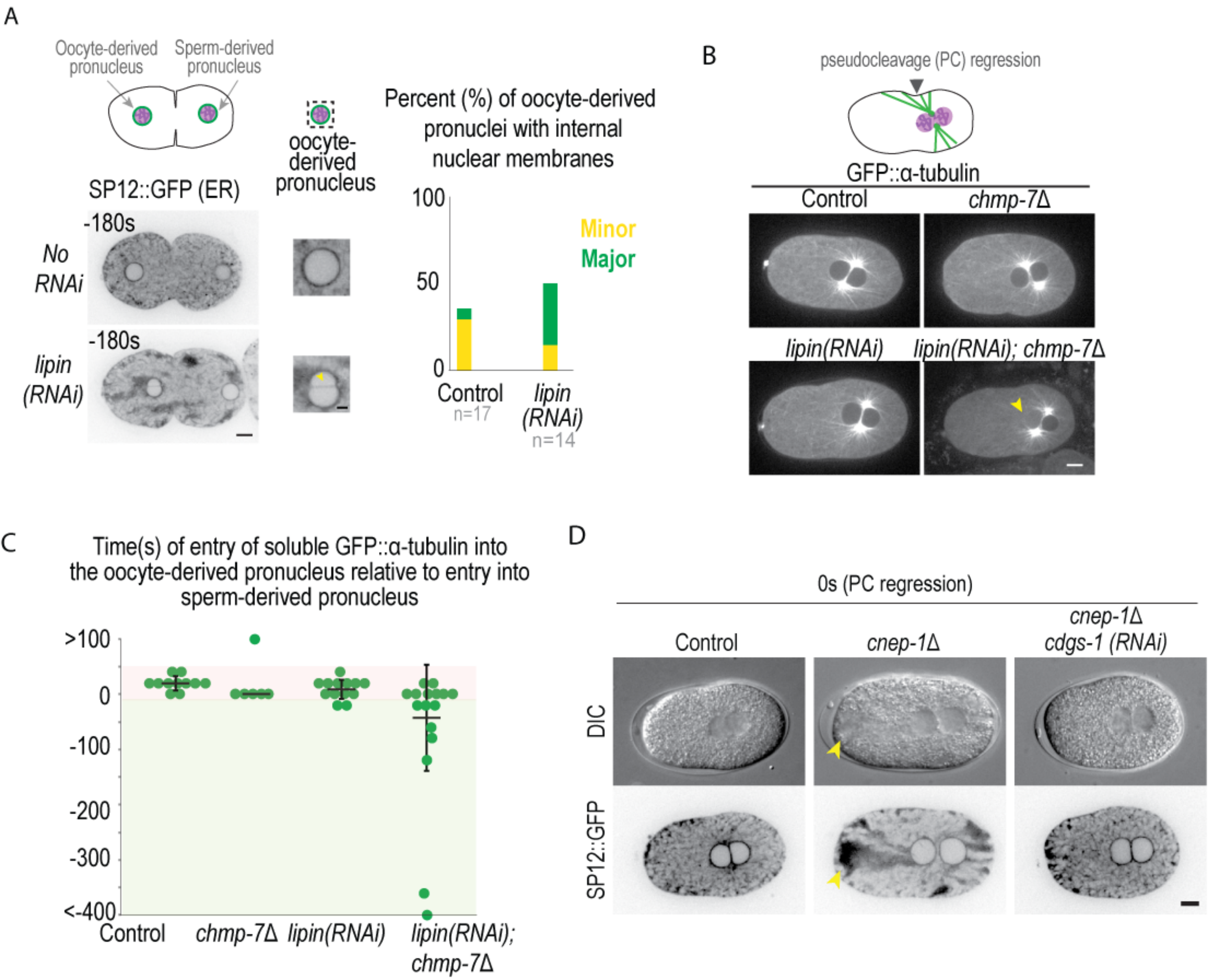
